# Drifting population dynamics with transient resets characterize sensorimotor transformation in the monkey superior colliculus

**DOI:** 10.1101/2023.01.03.522634

**Authors:** Michelle R. Heusser, Uday K. Jagadisan, Neeraj J. Gandhi

**Affiliations:** Departments of Bioengineering, University of Pittsburgh, Pittsburgh, PA, USA; Neuroscience, University of Pittsburgh, Pittsburgh, PA, USA; Center for the Neural Basis of Cognition, University of Pittsburgh, Pittsburgh, PA, USA

**Keywords:** sensorimotor transformation, population activity, superior colliculus, state-space framework, dimensionality reduction, saccade initiation, delayed saccade task

## Abstract

To produce goal-directed eye movements known as saccades, we must channel sensory input from our environment through a process known as sensorimotor transformation. The behavioral output of this phenomenon (an accurate eye movement) is straightforward, but the coordinated activity of neurons underlying its dynamics is not well understood. We searched for a neural correlate of sensorimotor transformation in the activity patterns of simultaneously recorded neurons in the superior colliculus (SC) of three male rhesus monkeys performing a visually guided, delayed saccade task. Neurons in the intermediate layers produce a burst of spikes both following the appearance of a visual (sensory) stimulus and preceding an eye movement command, but many also exhibit a sustained activity level during the intervening time (“delay period”). This sustained activity could be representative of visual processing or motor preparation, along with countless cognitive processes. Using a novel measure we call the Visuomotor Proximity Index (VMPI), we pitted visual and motor signals against each other by measuring the degree to which each session’s population activity (as summarized in a low-dimensional framework) could be considered more visual-like or more motor-like. The analysis highlighted two salient features of sensorimotor transformation. One, population activity on average drifted systematically toward a motor-like representation and intermittently reverted to a visual-like representation following a microsaccade. Two, activity patterns that drift to a stronger motor-like representation by the end of the delay period may enable a more rapid initiation of a saccade, substantiating the idea that this movement initiation mechanism is conserved across motor systems.

## INTRODUCTION

Sensorimotor transformation is a framework by which our brains process sensory input and produce a motor command. Its functionality is easily appreciated in the oculomotor system – when we see an object in our periphery, we can promptly direct our line of sight to that target. However, at what times are the neural populations representing the presence of a visual target through their coordinated activity? At what times are they collectively producing a signal that more closely resembles a motor command? And how does the population response transition from sensory to motor representations? The superior colliculus (SC) is a midbrain structure crucial for sensorimotor transformation (Basso and May, 2017; Cooper and McPeek, 2021; Gandhi and Katnani, 2011; Sajad et al., 2020; Wurtz and Optican, 1994). Neurons in its deeper layers emit strong bursts of activity both when a visual stimulus appears as well as when a high-velocity eye movement, known as a saccade, is generated to redirect gaze toward that object of interest. These putative “visual” and “motor” bursts are well characterized, but the time course of transforming visual stimulus-related information into a motor command is not understood to nearly the same degree. To this end, we searched for a neural correlate of sensorimotor transformation in small populations of SC neurons by characterizing the “visual-like” or “motor-like” pattern of activity during the intervening period of time between the visual and motor bursts while three male rhesus monkeys (*Macaca mulatta*) performed a visually guided delayed saccade task. This paradigm temporally separates the visual from the motor epoch through a “delay period” and has been previously employed in countless studies of cognition, sensation, and motor behavior.

If SC neural populations encode features independent of sensory information or movement preparation during the delay period (Kaufman et al., 2015), the activity patterns may lack a consistent trend in their visual- or motor-likeness across time. It could, for instance, meander within a distinct subspace (Figure 1A; Ames et al., 2019). However, if the SC systematically mediates sensorimotor transformation during this period, then there are a few candidate dynamics we could expect to observe. First, activity patterns could oscillate between a visual-like and a motor-like signal, balancing two needs – retaining information about the sensory stimulus and preparing for a movement (Figure 1B). This is akin to the idea of vacillating between population representations for two movements or stimuli (Caruso et al., 2018a; Dekleva et al., 2018; Kaufman et al., 2015; Rich and Wallis, 2016). Second, activity patterns could exhibit a discrete switch, or step, from one representation to the other (Figure 1C) (Latimer et al., 2015). Last, we could observe a smooth evolution in population activity patterns toward a motor-like signal, with a slow drift in representation between the times of sensation and action (Figure 1D). This last hypothesis is consistent with reports from single unit studies of sensorimotor integration in the oculomotor system. These studies have focused on reference frame transformations, with the objective of determining whether the temporally evolving neural activity better represents stimulus location or movement amplitude. The general result is that immediately after stimulus presentation, the sensory response is encoded in the reference frame of the stimulus modality – oculocentric for vision and craniocentric for audition. Just prior to the movement onset, the activity is best represented as a motor command in eye-centered coordinates or in a hybrid reference frame. In the intervening delay period, the average activity shows a slow and systematic transition from sensation to action representations, one which is sped up when no delay period is imposed. Such findings have been reported in the SC (Lee and Groh, 2012; Sajad et al., 2020; Sadeh et al., 2020), frontal eye fields (Caruso et al., 2018b; Sajad et al., 2016), parietal cortex (Buneo et al., 2002; Mullette-Gillman et al., 2005), and supplementary eye fields (Bharmauria et al., 2021).

**Figure 1.**
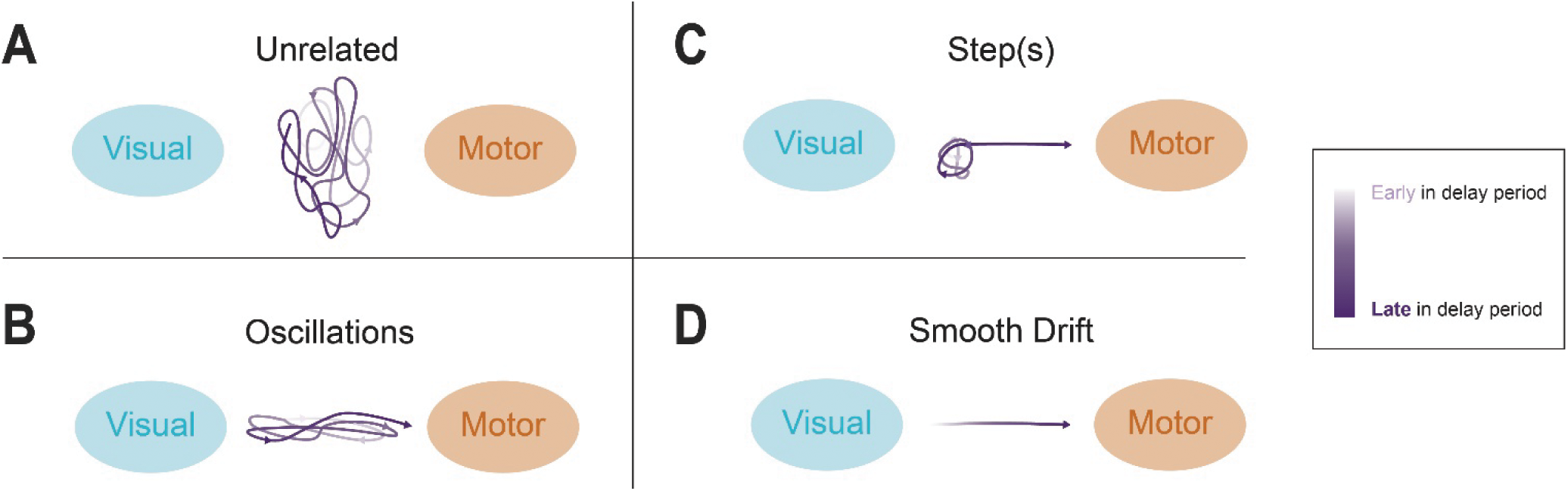
Schematic representations of four hypothetical delay activity neural trajectories. At one extreme, activity during the delay period is completely unrelated to sensorimotor transformation (**A**). Alternatively, the transition from sensation to action can display various features. For instance, the population activity can oscillate between the two representations (**B**), stay stagnant near the visual or some other subspace before abruptly stepping toward the motor subspace (**C**), or drift gradually from visual to motor representations during the delay period (**D**). The intensity of the purple color denotes the time course of a putative delay period.

To characterize the shared activity patterns of neural populations, we turned to a classic machine learning method known as dimensionality reduction. This family of algorithms has been utilized to investigate the dynamics of neural population activity underlying cognitive or behavioral processes such as stimulus encoding (e.g., Cowley et al., 2016), decision making (e.g., Aoi et al., 2020), and movement execution (e.g., Churchland et al., 2006). Such techniques transform the activity across the population into a state-space dynamical systems framework, where the pattern at any given moment can be represented as a linear combination of the activity of individual neurons. This methodology offers a noise-reduced, better-visualizable trajectory of activity across consecutive time points (Cunningham and Yu, 2014). Although these methods have already been readily accepted and widely utilized in other fields, they have also been employed sporadically in the oculomotor domain (Heusser et al., 2022; Jagadisan and Gandhi, 2022; Smalianchuk and Gandhi, 2022). In this study we aimed to enhance our knowledge of an important phenomenon using this increasingly popular framework, thereby demonstrating its applicability to the oculomotor system.

Here, we employed a dimensionality reduction algorithm called Gaussian Process Factor Analysis (GPFA, Yu et al., 2009), to characterize the time course of population-level representations as they relate to vision and saccadic eye movement. We then used a linear discriminant analysis (LDA) classifier to determine if the “subspaces” formed by collective, low-dimensional activity patterns during the visual and motor epochs were distinguishable from each other and found that for the bulk of neural populations, this was indeed the case. Exploiting this separability, we next computed the similarity of the activity patterns throughout the delay period to either the visual or the motor subspace through a Visuomotor Proximity Index (VMPI; based on the proximity index in Dekleva et al., 2018). Salient features of sensorimotor transformation emerged through this analysis. When looking across repetitions of the task, activity patterns exhibited a slow, systematic drift from a visual to a motor-like pattern. Remarkably, whenever a microsaccade occurred during the delay period, the population activity pattern transiently deviated to a visual-like representation before rapidly returning to the original trajectory. Finally, we tested an existing theory of arm movement generation known as the “initial condition hypothesis” (Afshar et al., 2011) and found that the state-space position of the activity on a given trial was correlated with the eventual saccadic reaction time, a relationship that emerged even hundreds of milliseconds before the cue to initiate a movement. These findings collectively clarify the SC’s role in sensorimotor transformation through both a network-level analysis of neural activity across sensorimotor epochs as well as a direct investigation of the relationship between this intermediate activity and behavior.

## RESULTS

Our objective was to characterize the delay period activity as exhibiting a visual- or motor-like signal and explore the relationship between this activity and the motor behavior to better understand the neural correlates of sensorimotor transformation in the SC. Neural activity from small populations of neurons was recorded simultaneously with multi-contact laminar probes traversing the dorsoventral axis of the SC as rhesus monkeys performed a visually guided, delayed saccade task (Figure 2). Data were examined from 27 recording sessions and limited to the subset of trials for which the visual stimulus and subsequent saccade were directed near the center of the response field. For electrode penetrations orthogonal to the SC surface, as we used here, all neurons recorded in a single session have comparable response fields, and care was taken to drive the electrode to the intermediate layers to capture mostly visuomotor neurons (i.e., those having both visual- and a motor-related increases in activity). A 24-channel laminar electrode was used for 15 of the sessions and a 16-channel electrode for the other 12 sessions. Spike sorting was performed and resulted in 12.3 (±3.3, range [7,19]) neurons per track, or population, on average. Across all 27 sessions, a total of 331 neurons were recorded.

**Figure 2.**
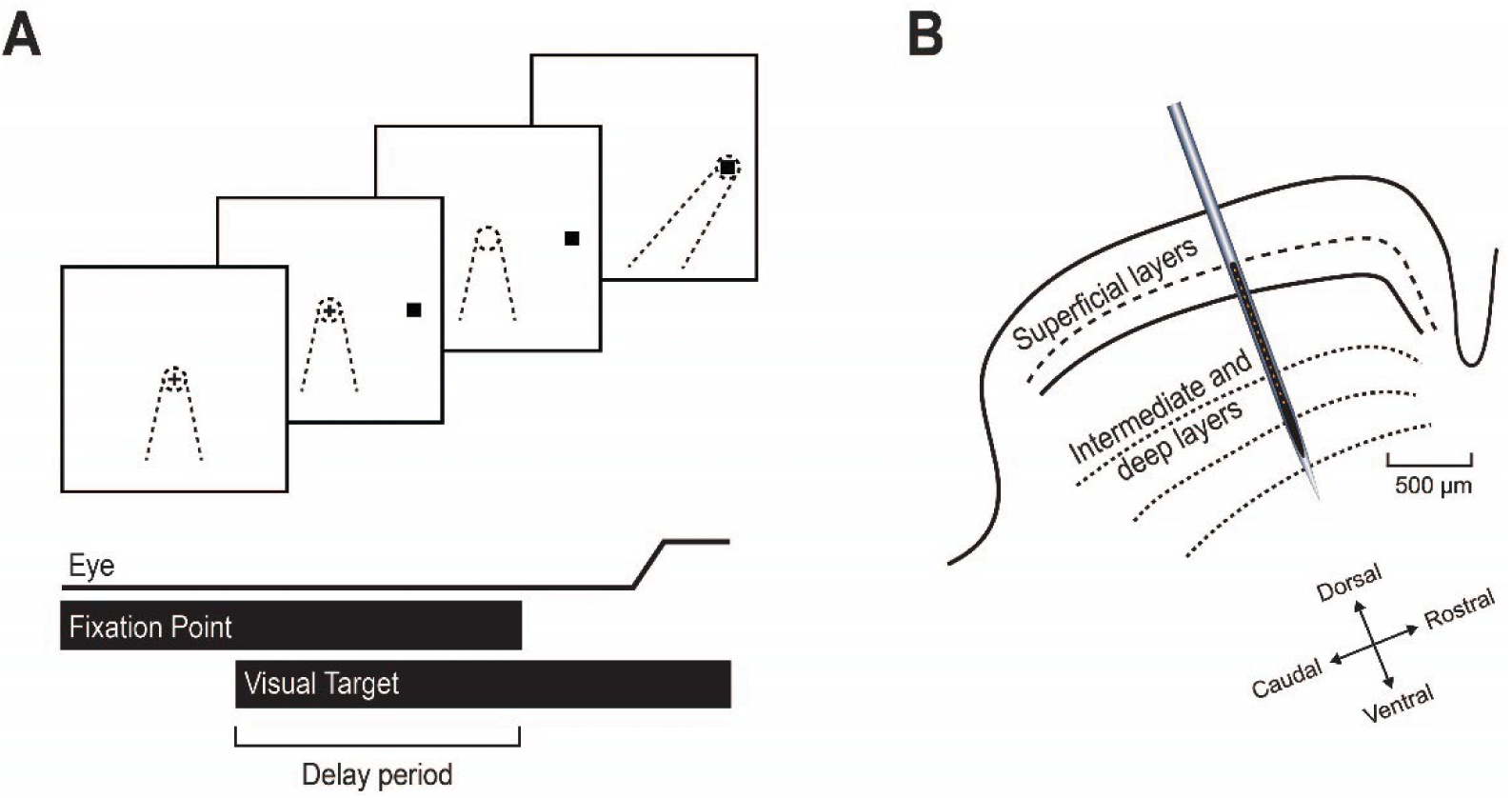
Schematic representation of behavioral task and neurophysiological preparation used in this study. **A.** Standard event sequence in the visually guided, delayed saccade task. Top – The animal fixates on a central point (shown as a plus sign in this illustration). A target (black square) appears in the periphery for a variable “delay period.” The dynamics of sensorimotor transformation were analyzed during much of the delay period. The disappearance of the fixation point acts as the go cue to generate a saccade toward the target. Center of gaze is depicted in all snapshots as a dotted cone. Bottom – A typical timeline of key events in a single trial of this task. **B.** In each experiment, a linear multielectrode array with 16 or 24 recording contacts was acutely inserted orthogonal to the SC surface along the dorsoventral axis to obtain a neural population representation as monkeys performed the delayed saccade task. Figure adapted from Jagadisan & Gandhi, 2022.

### Visual and motor subspaces are separable

We started by plotting for all contacts the average spike density profiles aligned on target and saccade onsets. We also separated the delay period from the transient visual burst, as shown for one session in Figure 3A. We then applied GPFA (Yu et al., 2009) to compute the latent activity patterns during these epochs (Figure 3B; see Methods). For most datasets (23/27), including this one, the top 3 factors accounted for at least 95% percent of the variance in the spike density profiles (Figure 3D). Thus, we limited our analysis to 3 dimensions, which also facilitated visualization. Moreover, the low-dimensional activity can be illustrated in a three-dimensional state space, in which a single point denotes activity across the population taken from a 20 ms window from one trial (Figure 3C). This framework, on which we base our first set of analyses, allows an assessment of the regions, or “subspaces,” where the activity resides and evolves across the various epochs of the trial.

**Figure 3.**
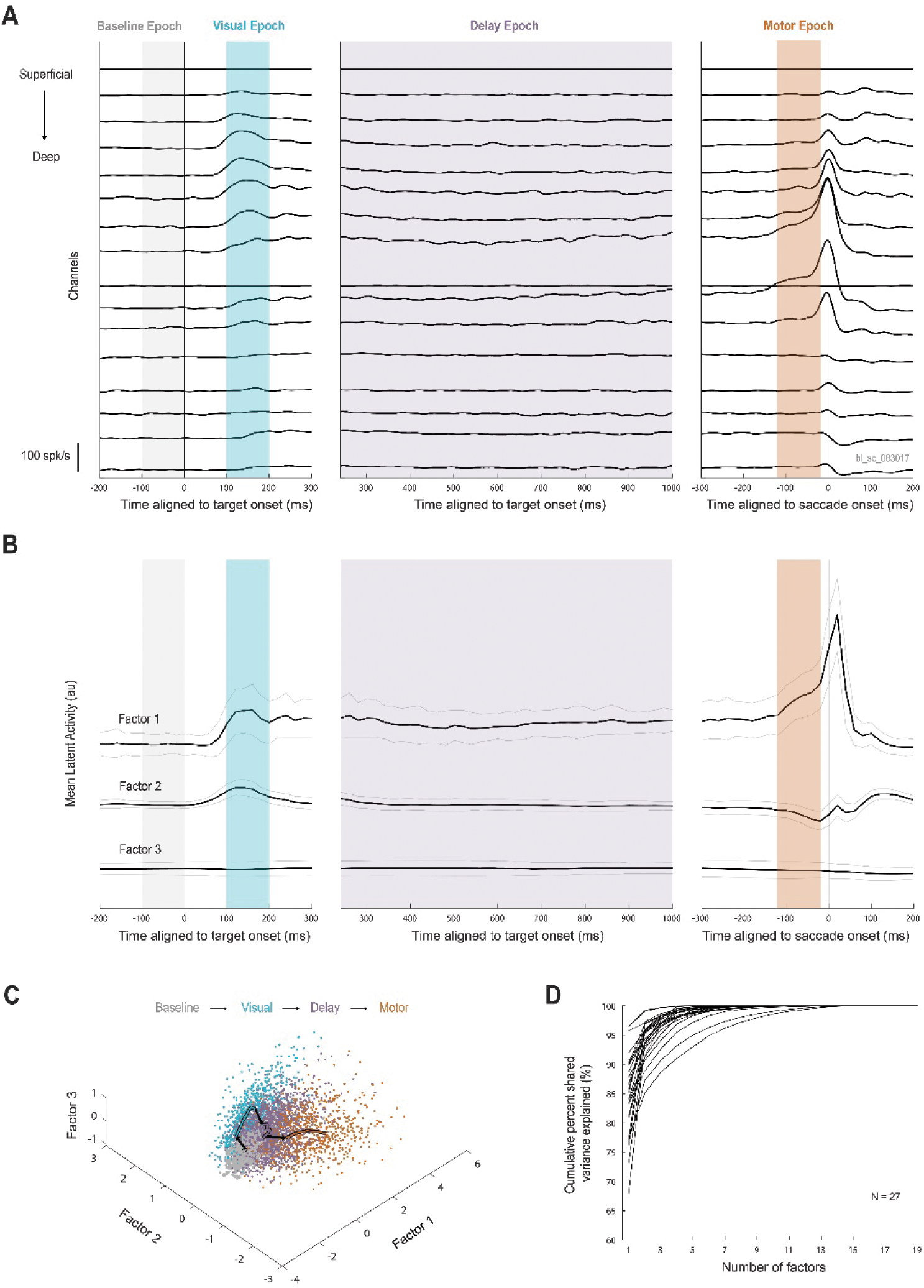
Analysis of neural population activity in a state space framework allows for an evaluation of subspace separability. **A.** Trial-averaged firing rates across electrode depth are shown for one example session. Multiunit activity on each of 16 channels is plotted aligned to target onset (left and middle) or saccade onset (right). Each epoch window is defined by a vertical rectangle (baseline in gray, visual in cyan, delay in purple, and motor in orange). Delay period activity was defined to start 240 ms after target onset, by which time the transient visual response had subsided. **B.** Latent population activity after dimensionality reduction using Gaussian Process Factor Analysis (GPFA) for the same example session after spike sorting into 12 single units. In each of the three panels, each trace is the trial-averaged (± one standard deviation) latent activity magnitude, plotted using the same alignment and epoch definitions as in (A). **C.** Latent activity represented in state space for the same example session. Latent activity during each of the four colored epochs are plotted as three-dimensional data points (each 20 ms bin has a magnitude along Factors 1, 2, and 3). Each dot represents the summary of the population activity pattern in a single 20 ms window; thus, a single trial contributes multiple points, even within the same epoch. The trial averaged trajectory in each epoch is layered above the individual points as a colored trace with black arrows denoting the direction of travel throughout the task timeline. **D.** Amount of covariability across neurons explained by lower-dimensional models compared to the full GPFA model. Each session is represented by a single trace. The majority of sessions have a high amount of shared variance explained by only one to three factors; thus, we retain the first three latent dimensions for each session.

In order to justify our comparison of delay period activity against two sets of activity, we first need to demonstrate that the activity patterns produced during the visual and motor epochs are distinct. Figure 4A-C shows the separability of the visual (100 to 200 ms after target onset) and motor (120 to 20 ms before saccade onset) latent activity patterns for three example datasets. By eye, the two subspaces are highly separable for these populations, and this separability was confirmed using linear discriminant analysis classification (Figure 4D). Across sessions, the mean classification accuracy was 82.7% (±11.3%), significantly above chance level of 50% (one-tailed t-test). This accuracy was not significantly different when considering multiunit activity (i.e., prior to spike sorting, see Supplementary Figure 1). Our subsequent analyses required a high level of separability between the visual and motor subspaces. We used a minimum classifier accuracy of 70% as our cutoff criterion, which reduced our yield to 22 datasets.

**Figure 4.**
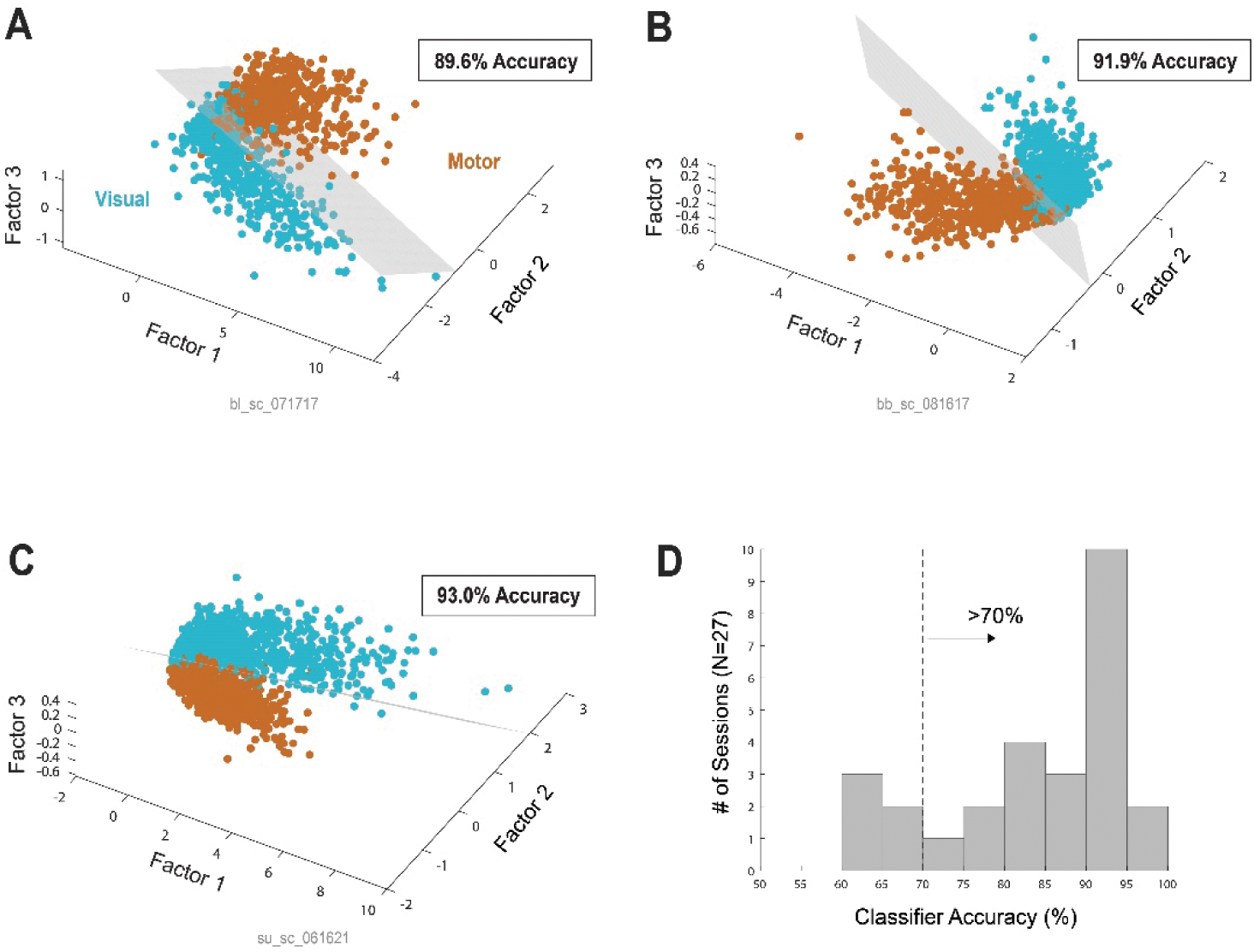
Visual and motor activity are linearly separable in state space for SC neural populations. **A.** Subspaces formed by latent activity patterns during the visual (cyan) and motor (orange) epochs for one example session. Time windows used for both epochs and a description of each data point are described in Figure 3. The gray shade indicates the plane of maximal separability as determined by linear discriminant analysis (LDA). The cross validated classification accuracy of visual and motor points for this example session is also reported. **B-C.** Same as in (A) but for two additional example sessions. **D.** Histogram of linear discriminant classifier cross-validated accuracy in distinguishing visual from motor patterns across all 27 sessions. Only sessions with accuracy values of 70% or better were used in analyses that assume high separation between visual and motor subspaces, leaving 22 sessions for future analyses.

### A gradual evolution from visual to motor subspace occurs during sensorimotor transformation

Once we established that visual and motor activity subspaces are separable, we wanted to examine the evolution of latent activity patterns throughout the delay period to determine whether there is a consistent trend in the activity pattern from a visual-like representation to a motor-like representation. We utilized a Visuomotor Proximity Index measure (VMPI, see Methods) to compute the similarity of the population activity pattern during small windows of time to the visual and motor subspaces (Figure 5). As expected, the VMPI is very close to the visual subspace during the visual epoch (cyan shade), to the motor subspace during the movement epoch (orange shade), and in-between during the delay period (purple shade).

**Figure 5.**
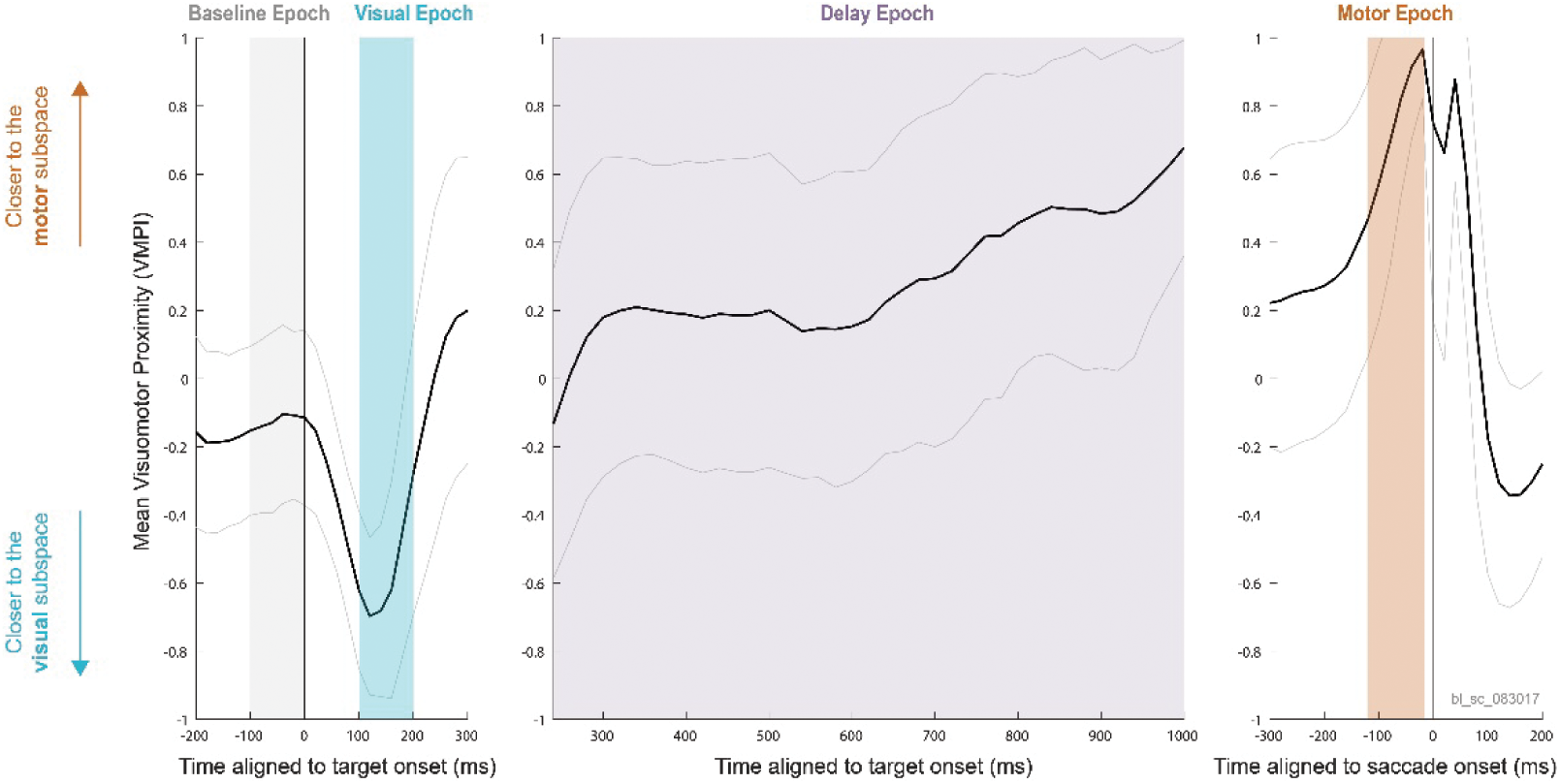
A Visuomotor Proximity Index (VMPI) can characterize the evolution of sensorimotor transformation. Mean (± one standard deviation) VMPI values across trials for the same example session as Figure 3. The three panels also obey the same alignment and color scheme. A value closer to +1 indicates similarity of activity to a motor pattern and a value closer to -1 indicates that activity is more similar to the activity pattern produced during the visual epoch. VMPI values are by design limited to the range [-1,1]. The evolution of VMPI across the delay period (purple shaded rectangle) give insights into the representations of SC population activity between the visual and motor epochs.

To evaluate the evolution of population activity, we plotted the VMPI during the delay period for all sessions before (Figure 6A) and after (Figure 6B) subtracting the mean across time bins for each session. These traces were then averaged to compute the session-averaged VMPI shown in Figure 6C. This trace reveals a slow, ramp-like progression of activity toward a motor-like representation as time in the delay period progresses. The mostly monotonic trend was highly statistically significant (p<0.001, Mann-Kendall test), suggesting that the neural activity pattern slowly and systematically drifts toward a motor-like representation. The evolution of VMPI was highly variable across trials, exhibiting unique but noisy dynamics on individual trials (see two example sessions in Supplementary Figure 2). Therefore, this monotonic trend only became evident after trial-averaging, as reported previously with single unit studies (e.g., Lee and Groh, 2012; Sajad et al., 2020; Sadeh et al., 2020). We also evaluated the time course of VMPI with the data aligned on the go cue (Figure 6D-F) and observed a similar but not as robust a trend (Figure 6F). This is not unexpected for data pooled over various delay periods for a sensorimotor transformation process that begins soon after target onset.

**Figure 6.**
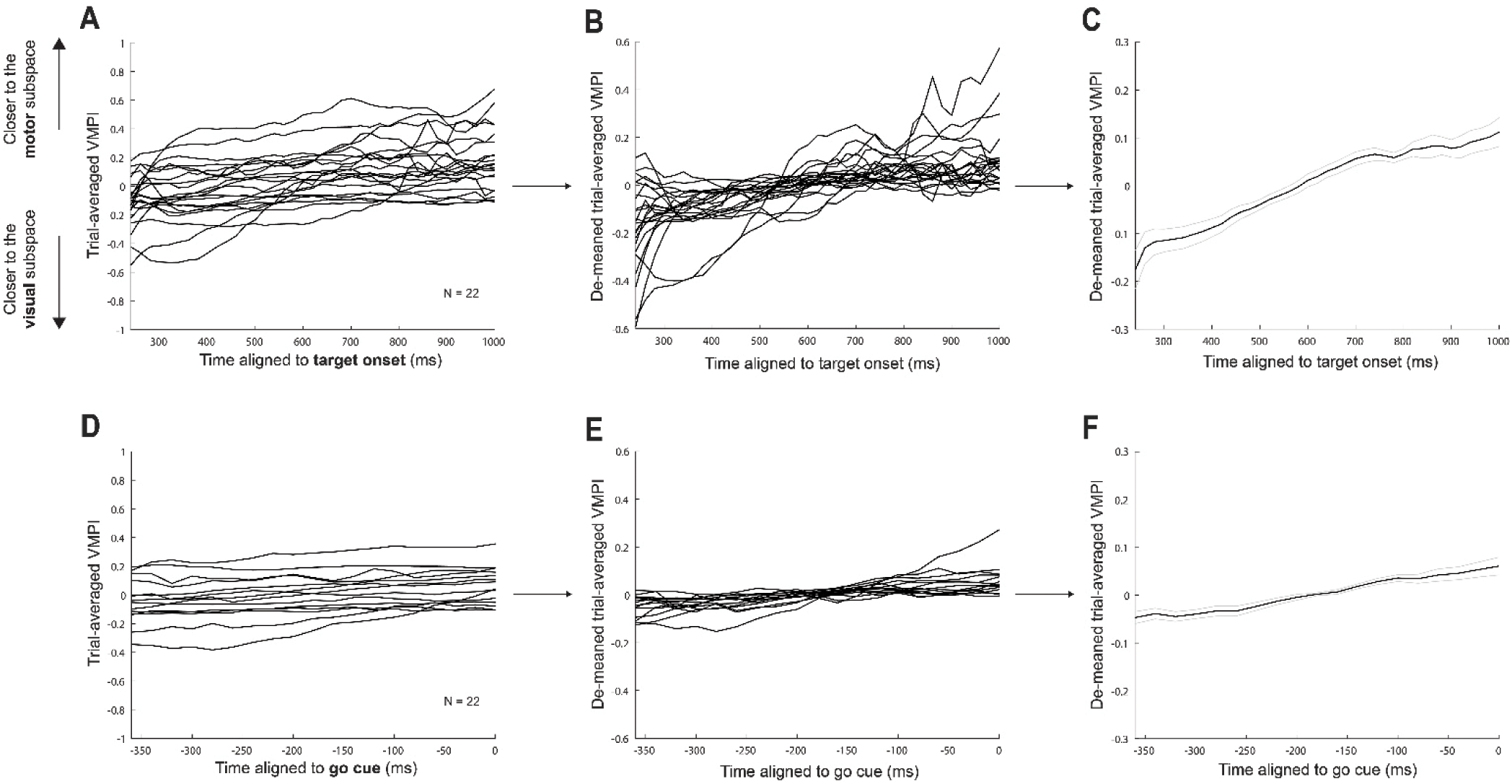
Trial-averaged neural activity slowly drifts from a visual to a motor-like pattern. **A.** Trial-averaged Visuomotor Proximity Index (VMPI) value across the delay period for the 22 sessions with >70% accurate separability between the visual and motor clusters. Each session is represented as a single trace. **B.** Same as in (A) but with VMPI traces de-meaned for each session separately. This allows us to average the VMPI traces across sessions to generate an across-session representation of the evolution of activity from the visual to motor subspace throughout the delay period, as shown in panel (**C**). There is a largely monotonic, highly statistically significant trend (p<0.001, Mann-Kendall test, based on raw traces in (A)) from a more visual to a more motor-like pattern throughout time in the delay period, indicative of a sensorimotor transformation signature. Note the progressively narrowed y axis limits across the three panels. **D-F.** Same as in A-C, respectively, but with average VMPI traces aligned to each trial’s go cue (equivalently, the end of the delay period). A similar though not as robust a trend in VMPI was observed for activity aligned on go cue, potentially due to dilution of effect from inclusion of different delay periods. Note the time scale differences of the x axes between panels A-C and D-F.

### Microsaccades transiently revert delay period activity toward visual subspace

Microsaccades are rapid eye movements with kinematics resembling those of larger saccades but characterized by their small magnitude (less than ∼2 degrees) and are frequently observed during fixational periods, including the delay period studied here (e.g., Hafed et al., 2015; Jagadisan & Gandhi, 2016; Peel et al., 2016). Therefore, we questioned if microsaccades that occurred during the delay period produced perturbations in the neural activity pattern that impacted the monotonic trend observed in Figure 6C. Microsaccades were detected offline (see Methods), and for sessions with a sufficient number of trials with microsaccades (N=14), trials in which at least one microsaccade occurred at any point during the delay period were separated from trials without microsaccades.

Interestingly, when we aligned the VMPI value on microsaccade onset (microsaccade trials) or a semi random delay period time (non-microsaccade trials, see Methods), we found a large and consistent effect, as illustrated in Figure 7A for an individual session’s data and Figure 7B as an across sessions average. Beginning approximately 50 ms after microsaccade onset, the VMPI deviated toward the visual subspace, indicating a pronounced shift toward a visual-like representation. This aligns well with the idea that microsaccades serve to refresh information about the visual stimulus in the SC by jittering the target location on the retina (Khademi et al., 2020). Even more compelling is the observation that the dynamical system compensates for the perturbation and returns the population activity to the normal (non-microsaccade trial) levels within roughly 100 milliseconds (Figure 7B).

**Figure 7.**
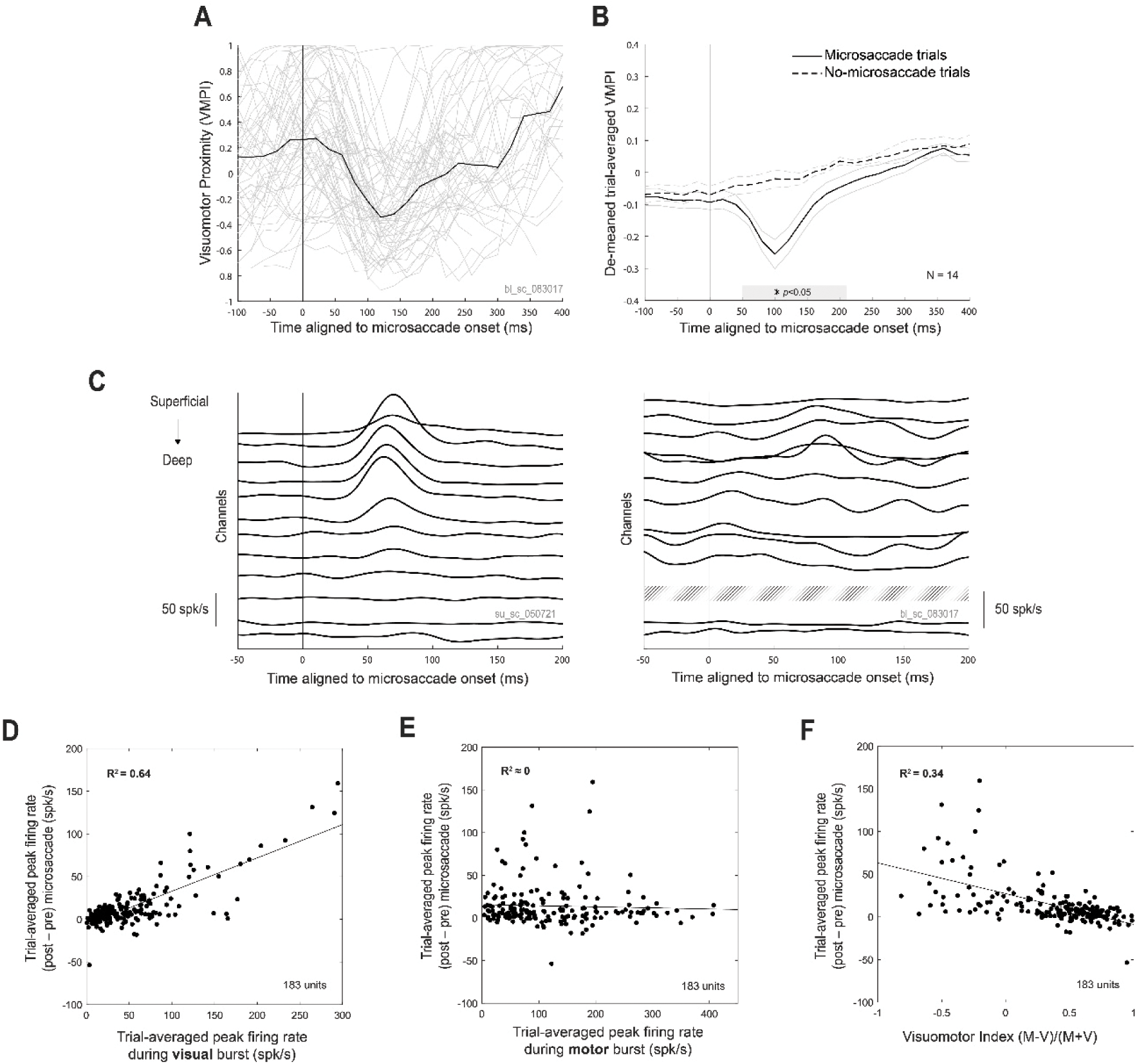
Single-trial population neural activity transiently reverts towards a visual pattern after a microsaccade. **A.** VMPI values on individual trials (gray) of an example session (same as Figure 3 and Figure 5) in which one or more microsaccades were detected during the delay period, with the median trace shown in black. VMPI values are aligned to microsaccade onset time, regardless of the absolute time in the delay period the microsaccade occurred. On average, the VMPI value dips toward a visual-like pattern ∼50 ms following a microsaccade. **B.** Session-averaged, de-meaned VMPI values (mean in black ± one standard error of the mean in gray, as in Figure 6C) aligned to microsaccade onset (solid lines) or pseudo-microsaccade onset on non-microsaccade trials (dashed lines, see Methods). The population activity pattern significantly and transiently deviates from the pattern produced on non-microsaccade trials starting around 50 ms and returns to match the pattern produced on non-microsaccade trials by 170 ms (Wilcoxon rank sum test, p<0.05). **C.** Trial-averaged firing rates of individual neurons aligned to microsaccade onset for two example sessions. Each unit is offset vertically based on the channel on which it was recorded (hence, the example session on the right has two units shown on one channel and no units shown on another, as indicated by diagonal shading). For many sessions, the superficial neurons burst following a microsaccade, although in some sessions – such as the example session on the right – it is difficult to appreciate any activity change following a microsaccade when looking at individual neuron firing rates. Still, even for these sessions, there is a population-level transient shift in representation toward a visual-like pattern (see same example session in (A)). **D.** Neurons with a visual response exhibit increased activity levels following a microsaccade. Each point represents a single neuron from a single session. **E-F.** Same setup as in (D) but for the relationship between peak activity around saccade onset (E) or visuomotor index (F) and the peak firing rate following a microsaccade. There is no correlation between the level of motor related and microsaccade-related firing rates, and there is a lower correlation between the relative visual and motor properties of neurons (using Visuomotor Index, VMI, as a proxy) and microsaccade-related activity than when only the strength of the visual burst is considered (i.e., D).

We decided to “zoom in” and see if this visual-like signature following a microsaccade was observable not only at the population level, but at the single neuron level. Figure 7C shows the trial-averaged firing rates of individual units aligned to microsaccade onset for two example sessions. For the session on the left, the more superficial units – those that typically exhibit a visual burst – clearly increase their firing rates following a microsaccade, consistent with the population-level analysis. This firing rate modulation of superficial units is nowhere near as pronounced for the example session on the right, yet the population-level measure we employed (i.e., VMPI) was able to easily pick up on a change in representation following a microsaccade (compare VMPI in Figure 7A with same session’s individual neuron dynamics in Figure 7C right panel). We probed this effect further by correlating across all neurons and sessions the post-microsaccade firing rate with the visual burst evoked when a stimulus is presented in the response field (Figure 7D, R^2^=0.64), the motor bursts generated for a saccade to that location (Figure 7E, R^2^ ≈0), and each neuron’s traditional visuomotor index (see Methods; Figure 7F, R^2^=0.34). The moderately strong relationship between visual activity and activity following a microsaccade confirms that the transient reset toward the visual subspace seen in Figure 7B in large part arises due to a strong transient increase in the activity of neurons in the population that have a visual burst.

### Sensorimotor transformation process is predictive of reaction time

We show above that on average, SC activity slowly drifts toward a motor-like representation throughout the delay period. This prompted us to ask if the magnitude of this drift on each individual trial was related to the animal’s ability to rapidly initiate an eye movement on that trial. We hypothesized that the higher the VMPI at a given time in the delay period (and therefore the larger the drift), the less time it would take the monkey to initiate the saccade after a cue to initiate movement (“go cue”) was given on that trial. Figure 8A shows the correlation coefficients between the VMPI value at time windows leading up to the animal’s go cue and the eventual saccadic reaction time (RT). At the time of the go cue to make an eye movement (time=0 ms, rightmost point of Figure 8A), the VMPI value was significantly correlated with the eventual RT, supporting the idea that the amount of drift on an individual trial by the end of the delay period is predictive of the animal’s ability to initiate a movement. This relationship held for all time points during the delay period leading up to the go cue (time range of -360 to 0 ms, Figure 8A). No other saccade metrics (i.e., amplitude, peak velocity, endpoint error) were found to be correlated with VMPI (Supplementary Figure 3A-B), nor might we expect them to be (Zhang et al., 2022). The presence or absence of microsaccades during the delay period did not qualitatively affect this relationship between VMPI and RT (Supplementary Figure 4).

**Figure 8.**
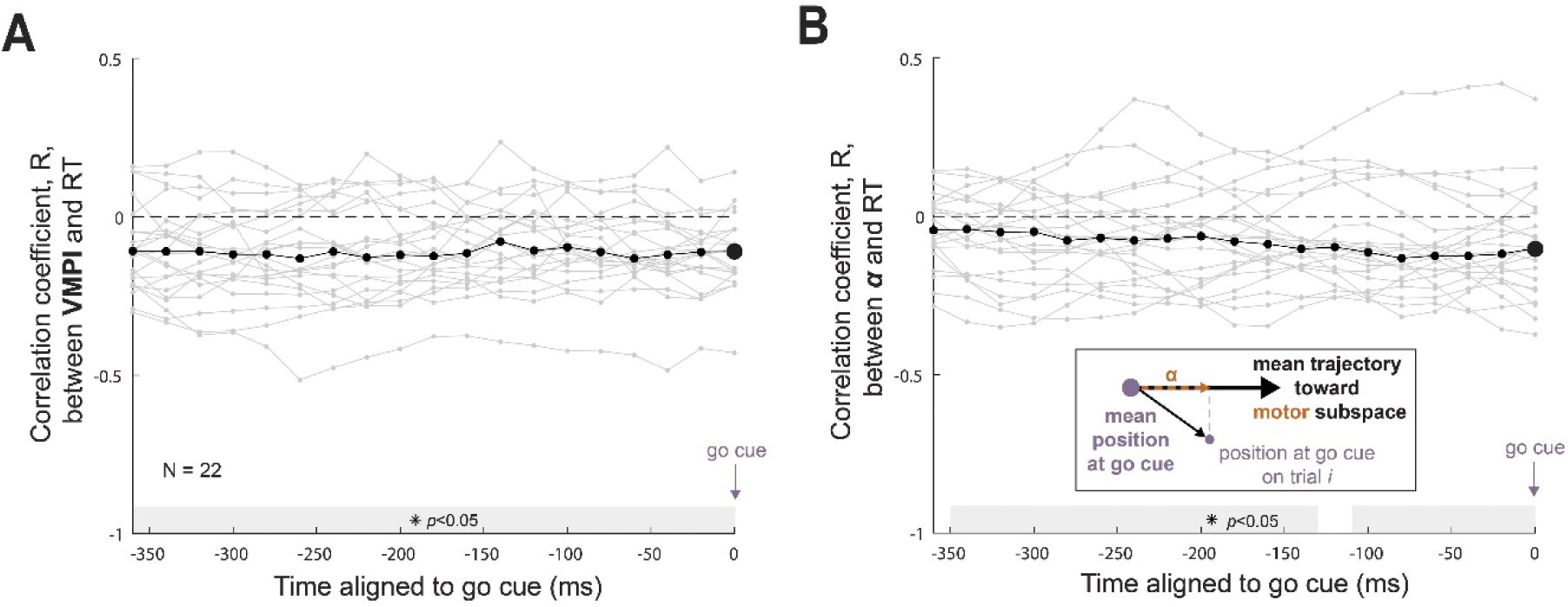
The single-trial state-space position of activity is correlated with that trial’s saccadic RT even long before the go cue. **A.** Across-trial correlation coefficient between the VMPI value in a single time bin relative to that trial’s go cue time and the eventual saccadic reaction time (RT) on that trial. Each gray trace represents the across-trial correlation coefficients for a single session, with the across-session median trace shown in black. Time bins in which the median correlation coefficients were significantly below zero (p<0.05, one-tailed Wilcoxon signed rank test) are shaded along the x axis in gray. Even long before the go cue, the state-space position of activity (as computed via VMPI) is correlated to a behavioral metric. **B.** Same as in (A) but for correlations between a single trial’s projection value *α* and that trial’s saccadic RT. Each projection value *α* represents the distance traveled along the trial-averaged neural trajectory toward the motor subspace by a certain time relative to the go cue on a given single trial *i*, with the method for finding *α* shown in the inset. Methodology was previously used in Afshar et al., 2011, and applied here to SC neural populations. This different method of computing state-space position of activity reveals a similar correlation to saccadic RTs, up to ∼350 ms before the go cue on average.

When aligning single-trial VMPI values to the beginning of the delay period (Supplementary Figure 3C), the position of the activity was not correlated to the eventual behavioral output, suggesting that the population representation likely drifts at relatively similar rates across trials. If so, then the drift would be bigger for the longer delay trials, which would be associated with faster reaction times. Indeed, the variable length of the delay period, in the context of our experimental paradigm, seems to account for the variable latency in the behavior (Supplementary Figure 5).

This finding conforms well to an existing theory of arm movement generation – the initial condition hypothesis (Afshar et al., 2011; Churchland et al., 2006) – in which the population activity pattern at the animal’s go cue informs the latency of the reach initiation on that trial. Critically, however, the brain areas relevant to reach initiation (e.g., primary motor cortex and dorsal premotor cortex) do not have significant responses to visual stimuli. Because of the strong visual bursts exhibited by SC neurons, we wanted to ensure that the relationship between population activity pattern and latency of saccade initiation remains even when disregarding the visual-likeness of the pattern during the delay period. Therefore, we decided to employ an additional methodology (mirroring that of Afshar et al., 2011) that only considers the position and trajectory of activity patterns relative to the patterns underlying *motor* output rather than relative to activity during both the visual *and* motor epochs (as is the case with the VMPI metric). In short, two vectors are created – one that extends from the mean activity position at the time of the go cue to some short time later (100 ms) and another that extends to the actual position at the time of the go cue (or before) on an individual trial (shown in Figure 8B inset). The projection value α obtained by projecting the latter vector onto the former gives you a magnitude that can be thought of as distance traveled toward the motor subspace by a certain time in the delay period.

When applying these methods to SC population activity, we found that the correlation values closely matched both those applied previously to premotor cortex activity and those obtained through our VMPI – RT correlation analysis. Across many populations, the median correlation between projection value at the go cue and RT across trials was small but significant (Figure 8B, data points at t=0 ms), similar to the results of the VMPI – RT correlation analysis (i.e., Figure 8A). Also comparable was the significant correlation between projection value α and eventual RT even long before the go cue (Figure 8B, data points leading up to t=0 ms). Therefore, this relationship between the population-level activity pattern and movement initiation latency holds even when only considering the motor-likeness of the pattern.

Last, we briefly shifted our focus to a single-neuron view to address whether the conclusions one might draw from our population-level analyses are consistent with those derived from the analysis of individual neurons. Indeed, we found that the firing rates of individual neurons when averaged over the entire length of the delay period were also correlated with single-trial reaction time (Supplementary Figure 6). Thus, a single-neuron view provides a congruent – albeit, more limited – interpretation of the relationship between SC activity during the delay period and subsequent motor behavior.

## DISCUSSION

In this study, we sought to understand whether population activity in the SC systematically transitions from a visual-like pattern to a motor-like pattern throughout the delay period. We labeled the transient burst representations that follow target onset and precede saccade onset as ’visual’ and ’motor’ subspaces, respectively, while remaining agnostic to their preferred coordinate system or reduced dimensional representation. We then sought to determine how, if at all, the population activity during the delay period transitioned between the two representations. This approach provides a more unsupervised yet still direct understanding of how SC populations encode these features. We found that on average, the activity pattern (as measured by the VMPI value) did indeed dynamically drift from a visual-like to a motor-like representation, consistent with the schematic shown in Figure 1D and with interpretations provided from previous studies on sensorimotor transformation in the SC (e.g., Lee and Groh, 2012; Sajad et al., 2020; Sadeh et al. 2020). Finally, we note that the gradual transition was only observed when averaging across trials.

Notably, following a microsaccade, the activity pattern was characterized by a transient reset to a visual-like representation (Figure 9A). In addition, the amount of drift exhibited by the population at times leading up to the animal’s go cue on individual trials was predictive of the latency at which a saccade could be initiated on that trial. On the other hand, neither the starting representation nor the amount of drift aligned to the beginning of the delay period were related to the eventual saccade latency. Together, these findings lead us to conclude that activity drifts at a relatively consistent rate and sets the animal up for a shorter or longer latency saccade based on the amount of time the activity has had to drift on a given trial (Figure 9B). This study therefore provides new insights into the neural dynamics expressed within the SC during the delay period of a widely used behavioral task.

**Figure 9.**
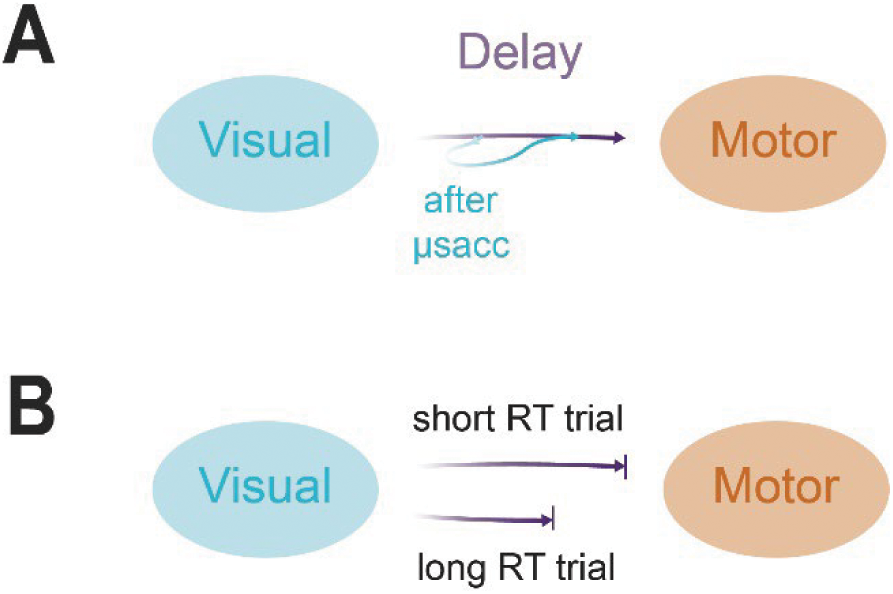
Schematics representing the evolution of SC population activity throughout the course of a standard sensorimotor task. **A.** Characteristic time course of population activity during the delay period. On average, population activity slowly becomes less visual-like (visual subspace shown as a cyan ellipse) and more motor-like (orange ellipse) throughout time in the delay period (purple line). Following a microsaccade (μsacc), when it occurs, population activity transiently and strongly deviates its trajectory toward the visual subspace (cyan line) before returning to the characteristic route toward the motor subspace. **B.** Depiction of the population activity trajectories on microsaccade-absent trials with short (top) and long (bottom) saccadic reaction times. Activity during the delay period likely evolves at a comparable rate across trials, at least under the experimental conditions we used. However, the activity can continue its slow drift toward the motor subspace for a longer time on trials with longer delay period lengths (top) than on those with shorter delay period lengths (bottom), thereby only necessitating a limited distance to travel to the motor subspace once the cue to initiate the behavior is given (vertical tick mark).

In one study, subpopulations of cortical oculomotor neurons categorized based on their relative firing rates during the visual and motor epochs were shown to exhibit unique temporal dynamics of this reference frame transformation, in contrast to the smooth and gradual transition observed when treating all neurons as a single population (Sajad et al., 2016). The VMPI measure we use implicitly takes into account the activities of all neural subtypes (i.e., visual, visuomotor, and motor) and produces one concise value, although the neurons recorded in our study were, by and large, visuomotor neurons. Research that specifically teases apart the individual contributions of neural subtypes to the time course of sensorimotor transformation may be required for a more complete understanding of the SC correlate of this behavioral phenomenon. Also not considered within our study are cognitive factors such as reward anticipation, arousal, and attention – factors which may in fact be multiplexed with encoded visual and motor signals present in the SC. Thus, the exact relationship of delay period SC activity patterns to phenomena other than visual processing and movement initiation is ripe for future investigations.

To our knowledge, only one prior study of oculomotor neural populations has taken a true dynamical systems approach to understanding the time-varying relationship of neural patterns to eventual movement metrics (e.g., RT). Khanna et al. (2019) performed GPFA on the spiking profiles of frontal eye fields (FEF) populations and found that for many latent factors, the activity values throughout the entire fixed-length delay period were statistically significantly correlated with single-trial RT. This result, although overall consistent with the present findings, does not directly consider the visual component of the activity; rather, it simply indicates that there is a general relationship between some pattern of population activity and movement metrics. Through our VMPI analysis, we made tangible the moment-by-moment representation of that population’s state-space configuration – specifically, whether it could be considered more similar to a visual pattern or to a motor pattern.

### Does SC delay activity resemble a preparatory signal?

The results shown in Figure 8A and 8B suggest that a relationship between the activity pattern and the reaction time (RT) of the eventual saccade is present long before permission is granted to initiate the movement. In other words, if the pattern of SC population activity drifts close to the motor subspace (i.e., has a high VMPI value) during the delay period of a given trial, the activity will take little time to evolve into a fully motor-like pattern after the go cue, resulting in a low-latency saccade. In the context of our task design, SC delay period activity seems to drift in representation at a relatively equal rate across trials (Supplementary Figure 3C). It is on trials with longer delays that the population activity has extra time to evolve, and therefore continues to drift toward a motor-like representation (equivalently, an increasing VMPI) proportional to the delay period length, resulting in proportionally fast reaction times (Supplementary Figure 5).

The observation that the rate of drift is unmodulated from one trial to another could be reflective of the animal’s internal model of the expected delay period length distribution. To extrapolate, if the delay period length was known by the animal to instead be constant, the activity pattern may still drift at slightly different rates from trial to trial, which we posit would constitute preparatory activity and serve as a mechanism for movement initiation. Regardless, the schematic shown in Figure 9 is inclusive of both schemas.

The consistent rate of sensorimotor evolution from trial to trial leads us to consider whether this drift might act as a self-timing mechanism, indicative of perceived time elapsed in the delay period. Suppose the monkey employs a strategy in which he begins to expect a delay period length somewhere in between the shortest and longest delays previously experienced (as observed in human studies; Jazayeri & Shadlen, 2010). If he consistently plans to initiate an eye movement after this expected delay period length, he may receive the benefit of more frequently successful trials (and consequently, more frequent and more rapid rewards) since he has optimized the timing of his saccade to match his expectation. Neurons in the macaque thalamus (Tanaka, 2007) and lateral intraparietal cortex (Leon and Shadlen, 2003) have been shown to encode perceived time intervals. Perhaps the drifting representation we observe in SC populations is another signature of task timing. Further experiments might explore this concept and its validity.

Although we consider SC activity during the delay period to be preparatory in the sense that it is related to the enhancement or hindrance of rapid saccade initiation following the go cue, it does not have “motor potential.” In the smooth pursuit system of the FEF, neural activity *was* found to have motor potential, with partially overlapping subpopulations contributing to both the preparation and execution of movement (Darlington and Lisberger, 2020). However, this does not seem to be the case for the SC, at least in the context of saccades. Previously, we have demonstrated that inhibition of the omnipause neurons during the delay period, which allows SC activity to travel to saccade-generating brainstem structures, is not sufficient to evoke a saccade (Gandhi and Bonadonna, 2005; Jagadisan and Gandhi, 2017). In addition, the lack of a burst and only a baseline-level “temporal stability” – that is, the consistency in the population activity pattern from one time point to another – enhance the argument against delay activity having motor potential (Uday K. Jagadisan and Gandhi, 2022). We have also observed that it is typically the neurons with strong visual bursts rather than strong motor bursts that have sustained activity during the delay period (Massot et al., 2019). Therefore, it stands to reason that preparatory signals are likely encoded in the SC in dimensions orthogonal to those during movement (e.g., orthogonal potent-null subspaces; Kaufman et al., 2014).

### The relationship between microsaccades and SC population activity patterns

Microsaccades produced during fixation serve to refresh the visual stimulus on the retina in order to combat a fading perception over time (Martinez-Conde et al., 2004). The neural circuit in SC suppresses vision during microsaccades (Hafed and Krauzlis, 2010), as it does during large amplitude saccades (Robinson and Wurtz, 1976). Following the movement, the nervous system responds to its visual environment by evoking activity in visually responsive neurons, although extra-retinal sources likely contribute as well. Indeed, we observed that microsaccades produced during the delay period consistently perturbed the sensation to action transition by transiently deviating SC population activity toward the visual subspace. The effect was strongest in the subset of neurons with a robust visual response. This modulation began roughly 50 ms after microsaccade onset and peaked another 50 ms later before rapidly meeting back up with the population activity patterns observed in non-microsaccade trials. Thus, the resurgence of visual activity likely reflects visual reafference following the movement (also see Khademi et al., 2020).

It is valuable to consider the various ways in which microsaccades generated during the delay period could have impacted the oculomotor system. For instance, the movement-related activity associated with microsaccade generation could have accelerated the overall SC output toward the motor subspace, resulting in reduced saccade latency – perhaps even triggering it prematurely before the go cue – and altered endpoint accuracy (Buonocore et al., 2021). Instead, we observed a rapid rebound and return of the system (VMPI trace, Figure 7B) to its original trajectory following a microsaccade. We interpret this reset to suggest that the gradual transition from a sensory to motor representation may be a network feature that is resistant to the effects of transient disruptions. As a whole, these observations lead us to conclude that microsaccades are a potential mechanism for engaging the network to produce a visual-like signal very similar to that elicited in response to the initial target appearance, but one that is compensated for quickly and robustly.

### Low-dimensional geometry of SC population activity and its skeletomotor counterparts

One of our objectives was to extend to the oculomotor system the dynamical systems perspective of motor control that has been studied extensively in the skeletomotor system (Gallego et al., 2017; Shenoy et al., 2013). Studies of arm reaching that use this framework have given rise to multiple hypotheses for mechanisms of movement initiation. One such schema is the “optimal subspace hypothesis” (Churchland et al., 2006), which propounds that there is an optimal set of population activity patterns that allow for the generation of a goal-directed movement. The initial condition hypothesis (Afshar et al., 2011) builds on this framework by postulating that trials in which patterns that have traveled closer to the motor subspace by the time of the animal’s go cue will have a faster reaction time (RT) than those in which the underlying neural activity has not traveled as far along the mean neural trajectory.

It might make sense for the oculomotor system to operate in a somewhat different manner than the skeletomotor system given the additional element – visual information – encoded within the SC and other oculomotor areas. However, even when considering this additional set of patterns exhibited by SC populations, we found that on trials in which population activity more closely resembles a motor-like pattern, the saccadic RT is significantly shorter (Figure 8A).

To establish a more direct comparison between SC activity and activity in its skeletomotor analogs, we also applied the methods of Afshar et al., 2011, to SC population activity during the delayed saccade task and found a comparable, significant correlation between the position of delay period activity and the saccadic RT (Figure 8B). Although the exact methodologies applied in Figures 8A and 8B are distinct, they address similar questions – primarily, is the similarity of neural activity during the delay period to motor activity related to the speed at which the movement can be initiated (i.e., eye movement or reaching movement RT). The findings reported in this study support the idea that the initial condition hypothesis is also valid for the oculomotor system.

The optimal number of dimensions needed to explain the across-trial shared variance of our acutely recorded neural populations was much lower than that described in studies using neural activity recorded in primary motor cortex (M1) or dorsal premotor cortex (PMd), for example (Churchland et al., 2010; Churchland and Shenoy, 2007). We conjecture that this is at least partially due to the homogeneity of each recorded population. As SC neurons are traditionally recorded along a dorsoventral axis, the topography of the SC yields populations in which each neuron has a similar response field, chiefly varying across electrode depth in the strength of their visual and motor bursts (Bourrelly et al., 2023; Massot et al., 2019). Cortical areas like M1 and PMd yield much more heterogenous populations with respect to the spatial locations preferentially encoded by each neuron, and the dynamics underlying behavior are typically studied after grouping trials with multiple reach directions. However, since our recorded SC populations vary not in their preferred spatial location but rather in their visual and motor signal strengths, we limited our analyses to a single saccade direction so that in reducing the dimensionality of the data, the variability between visual and motor patterns would be brought to the forefront.

## MATERIALS AND METHODS

### Subjects and surgical approach

Three adult male rhesus monkeys (*Macaca mulatta;* monkeys BL, BB, and SU) were used for this study. The experimental protocol was approved by the University of Pittsburgh Institutional Animal Care and Use Committee. Each animal underwent a sterile surgery under general anesthesia to implant a cylindrical recording chamber (Narishige) positioned above a craniotomy that allows access to the SC. A Teflon-coated, stainless-steel wire was also implanted on one eye in some animals. Surgical methods are described in more detail in Jagadisan & Gandhi, 2016.

### Visual stimuli and behavioral paradigm

Stimulus presentation and the animal’s behavior were under real-time control with a LabVIEW-based controller interface (Bryant and Gandhi, 2005). All stimuli were white squares, 4×4 pixels subtending approximately 0.5°, displayed against a dark grey background on a LED-backlit flat screen monitor. Eye position was recorded using the scleral search coil technique (CNC Engineering) or using an EyeLink 1000 eye tracker (SR Research), both sampled at 1 kHz.

Each monkey was trained to sit head-restrained in a primate chair and perform a standard delayed saccade task in a dimly lit room. To complete a successful trial of this task, the monkey fixated on a visual stimulus located in the center of the screen and maintained fixation while a visual target was presented in the animal’s periphery. After a variable delay period (600-1200 ms for monkeys BL and BB, and 700-1500 ms for monkey SU, weighted to maintain a flat anticipation function), the fixation point was extinguished, serving as the animal’s “go cue” to make a saccadic eye movement to the target. The animal had to make a saccade to the peripheral target within 460-800 ms and was required to maintain fixation on it for 300 ms to receive a liquid reward. The monkeys performed this task with high accuracy before recording sessions began. Thus, we limited our analyses to rewarded trials only.

### Electrophysiology and data pre-processing

On each recording session, a 16- or 24-channel linear microelectrode array (Plexon or AlphaOmega) was inserted orthogonal to the SC surface along the dorsoventral axis. Neurons recorded using this approach had similar preferred saccade vectors as determined by microstimulation (Massot et al., 2019). Care was taken to position the electrode in a way that maximized the yield of neurons exhibiting both visual and motor bursts, and thereby tended to be positioned in the deeper SC layers.

On each given trial, the target presented in the animal’s periphery could be located near the center of the response field of the recorded neurons (as determined by microstimulation) or in the diametrically opposite position with a 2:1 ratio of occurrence. Only correct trials in which the target was presented in the recorded neurons’ response field were included in analyses, and all analyses were performed separately for each neural population. Trials were further limited to only those with saccadic reaction times of greater than 100 ms to remove potential “cheat” trials. Unless otherwise specified, all analyses were performed using MATLAB 2019a (MathWorks) with custom code.

Spike times on each channel were first obtained offline using a voltage thresholding method. Each channel’s spiking activity was then manually sorted into single units before continuing with analyses (using MKsort, a spike-sorting user interface, Ripple Neuro), and the low-dimensional representations of population activity are quite similar (see Supplementary Figure 1). This is in line with previous work demonstrating that spike sorting has a negligible effect on the message of studies that focus on low dimensional dynamics of population neural activity (Trautmann et al., 2019). Therefore, throughout the text we present results from spike-sorted neural activity but think of units (neurons) and multiunits (channels) as interchangeable. A total of 27 sessions were obtained and included in this study.

### Dimensionality reduction

In order to analyze the spiking patterns across the entire population, we utilized a dimensionality reduction method called Gaussian-process factor analysis, or GPFA (Yu et al., 2009). In short, this method converts spiking activity from a neural population into a lower-dimensional continuous “neural trajectory,” where each dimension represents a weighted linear combination of neurons.

To perform GPFA, we used DataHigh (Cowley et al., 2013), a publicly available MATLAB code package for visualizing and reducing dimensionality of high-dimensional neural data. For a given session of laminar electrode data, all channels’ spike times were first converted into spike trains aligned on target onset. Spike counts were grouped into non-overlapping bins of 20 ms width. Each observation includes data from one trial of the delayed saccade task beginning 200 ms before target onset and continuing through 200 ms post-saccade. This matrix is of size N channels x T time bins for each trial, with the latter dimension having a variable length. The GPFA algorithm returns a set of latent activity values summarizing the activity pattern across the population for each trial (matrix of L latent dimensions x T time bins for each trial). A cross-validation procedure was performed to determine the optimal number of reduced dimensions. The optimal dimensionality for each found via cross-validation was typically low (one to three). A final dimensionality of three was chosen for the sake of consistency across sessions and for ease of visualization in a 3D state space, as described next.

### Defining subspaces and computing proximity

The term “subspace” does not have a widely agreed-upon definition; some groups call each factor returned by dimensionality reduction a subspace in which latent neural activity could be varying (e.g., Kaufman et al., 2014) while others define new axes and focus on the variable activity along that dimension (e.g., Kobak et al., 2016; Libby & Buschman, 2021) or project activity onto an axis, plane, or hyperplane contained within a higher-dimensional space (e.g., Aoi et al., 2020; Semedo et al., 2019). Here, we more loosely define a subspace as a distinct region in a low-dimensional state space occupied by neural activity during a specified condition (as in Churchland et al., 2006). The two main conditions here are “visual,” or 100 to 200 ms after target onset (around the time of the putative visual burst) and “motor,” or 120 ms to 20 ms before saccade onset (around the rising phase of the putative motor burst, and not including activity from after the latest time it is likely related to saccade initiation; see (Gandhi & Keller, 1999; Jagadisan & Gandhi, 2017; Miyashita & Hikosaka, 1996; Smalianchuk et al., 2018). A baseline condition was also defined, and this includes activity from 100 ms before target onset up to the time of target onset. These comprise the visual, motor and baseline subspaces, respectively.

For Figure 4, linear discriminant analysis (LDA) was performed to find the 2D projection that best separates visual and motor subspaces from each other. For each session, a two-class (“visual” and “motor” categories) linear discriminant classifier was trained and tested using a 10-fold cross-validation procedure. Only sessions for which the population activity was sufficiently separable between the visual and motor epochs (>70% classification accuracy rounded to the nearest integer, see Figure 4D) were further analyzed, leaving 22 sessions that met this criterion.

Given that the visual and motor activity (cyan and orange points, respectively) form distinct subspaces, we can use these as reference distributions against which to compare activity from time points throughout the course of each trial. We utilized a measure known as the proximity index, introduced by Dekleva et al., 2018. In short, the proximity index is a probabilistic measure that indicates the relative likelihood that a point of activity is closer in state space to a particular cluster than any other comparison cluster. For a single time bin of latent activity S, its proximity to the visual or motor cluster (VPI or MPI, respectively) is given by:

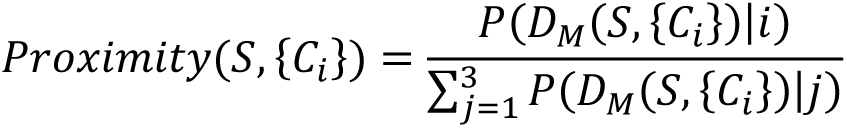

where {*C_i_*} is the cluster of latent activity points during one of three reference conditions, denoted by *i* and *j* (here: visual, motor, or baseline) and *D_M_*(*S*, {*C_i_*}) is the Mahalanobis distance between the point S and cluster {*C_i_*}. The VPI is formed when *i* =visual activity (and *j* =motor and baseline) and likewise, the MPI is formed when *i* =motor activity. The VPI and MPI are normalized to the range [0, 1].

Since visual and motor proximity indices must be computed separately and result in two yoked values, we defined a visuomotor proximity index (VMPI) that can range from −1 to +1 and gives the relative proximity value of a point of activity to either the visual (−1) or motor (+1) subspace:

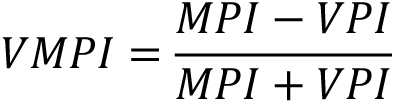

For Figure 5, the VMPI was computed for all non-overlapping 20 ms bins of latent activity throughout the time course of each trial. Activity from the baseline condition was treated as a third cluster to allow for the possibility of delay period activity existing in a completely different subspace than either the visual or motor subspaces, although proximities to this cluster are unimportant and hence not shown. It is also important to note that the absolute VMPI value ranges are inconsistent across populations (e.g., Figure 6A), but this does not matter for our study. Only the dynamics in the VMPI trace over the course of the delay period are of interest; thus, in Figure 6B-C and Figure 7B-C each population’s VMPI trace was mean-subtracted to allow for a better comparison of sensorimotor transformation across populations.

### Detecting microsaccades and aligning proximity to microsaccade onset

All microsaccades that occurred during the delay period of each trial were detected offline. A 20 ms moving average of the eye velocity was taken, and a speed threshold of 5 to 15 deg/s was applied depending on noise level in the eye position signal. Saccades greater than 2 degrees in amplitude were rejected. Individual trials were manually evaluated to confirm correct automatic detection. Sessions were included in the following analysis if there were at least 20 trials in both the “one or more microsaccades” and “no microsaccades” conditions. One monkey (BB) did not consistently produce microsaccades during the delay period, as we have reported previously (Jagadisan and Gandhi, 2016). Hence, we could only include data from one session for this monkey using the above criteria. Monkeys BL and SU had five and eight sessions that met the above criteria, respectively, for a total of 14 sessions included in this set of analyses.

To determine the effect of microsaccades on the population activity pattern, we aligned the VMPI to microsaccade onset for trials in which at least one microsaccade was detected during the delay period. As a control analysis, we also aligned the VMPI to a pseudo microsaccade onset time for trials in which there was no microsaccade detected. For each trial, this alignment time was created by selecting a random time from the distribution of microsaccade onset times in trials with a microsaccade (Figure 7A-B). As in Figure 6, each population’s trace was mean-subtracted to better compare trends across sessions.

### Computing relationship between population activity patterns and behavioral metrics

To examine whether the position in state space is related to the end behavior (i.e., saccade), we computed the correlation between the VMPI value at the animal’s go cue and the eventual saccadic reaction time (RT) on that trial. We also asked if the position of the activity even leading up to the go cue was correlated with the end behavior. For this analysis, we worked backwards to compute the correlation coefficient of every 20 ms bin of activity with the saccadic reaction times on their respective trials. Values were tested for significance using a Wilcoxon rank sum test.

We also employed a similar approach developed by Afshar et al., 2011, that only utilizes information about the putative motor subspace rather than the visual subspace. In this framework, one can ask whether the distance traveled along the mean neural trajectory at the end of the delay period (equivalently, at the time of the animal’s go cue), correlates with the saccadic RT on that trial. These methods have been described previously and were followed as closely as possible. In short, on a single trial, a vector of spike counts across the population starting at the time of go cue and going forward some short time in the future (dt=100 ms) is created. This vector is projected onto the vector created by mean values across all trials to obtain a projection value α (see Figure 8B inset). A correlation coefficient value between saccadic reaction time and this projection value was obtained for each session. To compute the correlation between activity prior to the go cue and the end behavior, we used the same value of dt (100 ms) but worked backwards to compute the median correlation coefficient of every 20 ms bin of activity with the RTs on their respective trials, as in the VMPI – RT correlation analysis. Of note, unlike all other results presented in this paper, this analysis was performed on spike sorted but not dimensionality-reduced neural activity for a more direct comparison of findings across brain areas.

## ACKNOWLEDGEMENTS

N.J.G. was supported by NIH R01 EY024831 and R01 EY022854. M.R.H. was supported by the ARCS Foundation, GAANN Fellowship Program (P200A150050), and Behavioral Brain Research Training Program (NIH T32 GM081760). The authors would like to thank Dr. Clara Bourrelly for data collection, Dr. Benjamin Cowley for thoughtful discussions, and Dr. Brian Dekleva for providing a template code for the proximity analysis.

## SUPPLEMENTARY FIGURES

**Supplementary Figure 1.**
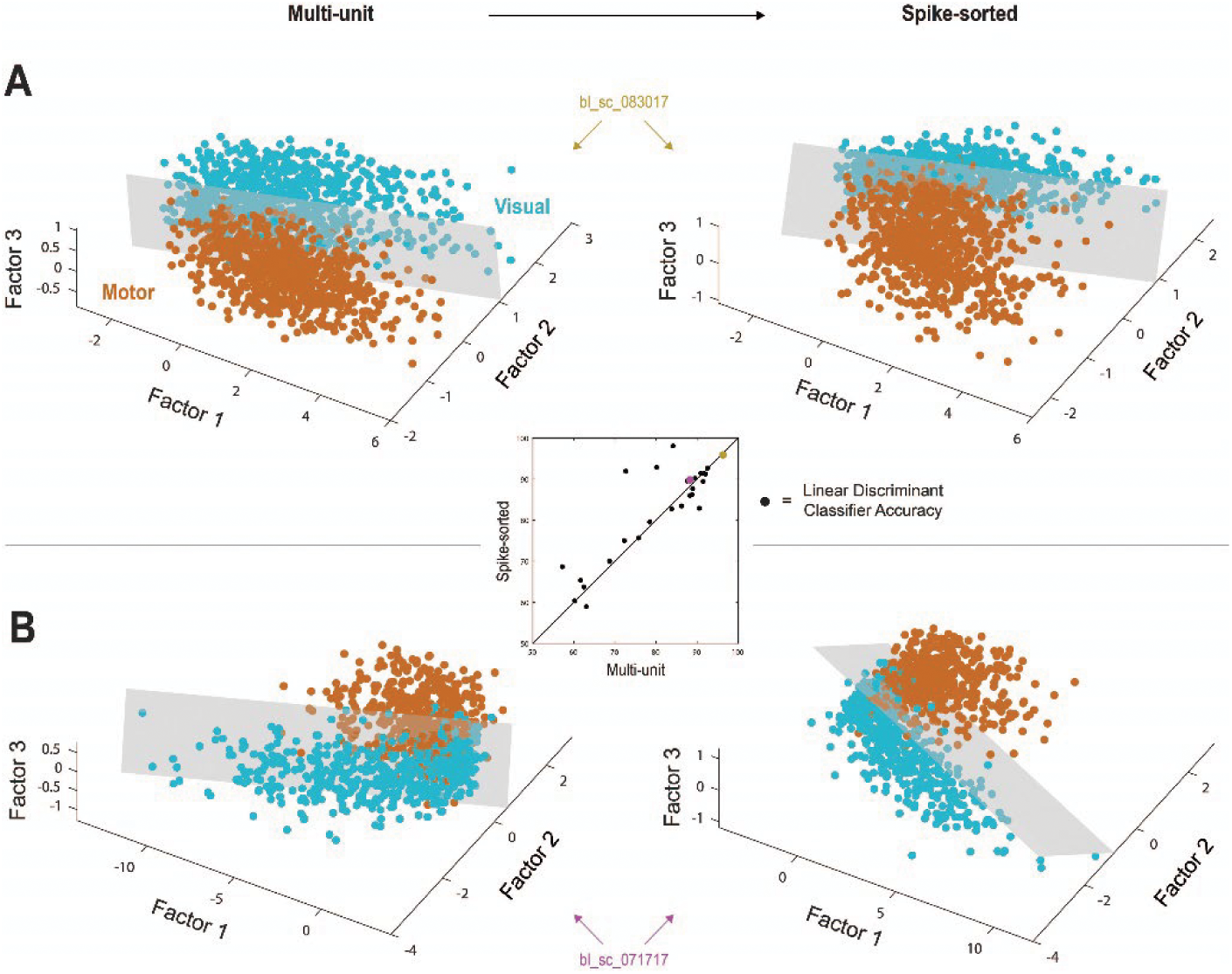
Spike-sorted and multiunit populations exhibit nearly identical activity patterns throughout the delayed saccade task. **A.** Low-dimensional representations of population activity patterns during the visual (cyan) and motor (orange) epochs for one example session before (left) and after (right) spike sorting. Although each dimension is not directly comparable across multiunit and spike-sorted populations, the subspaces formed by population activity in both cases are nearly identical. **B.** Same as in (A) but for a second example session. The subspaces formed by visual and motor activity before and after spike sorting have similar levels of separability. The exact position of each point of activity is unimportant to the comparison across epochs. **Inset.** A comparison of the visual and motor subspace separability obtained through LDA classification pre (x axis) and post (y axis) spike sorting for all 27 sessions. Accuracies for example sessions shown in (A) and (B) are colored in gold and magenta, respectively.

**Supplementary Figure 2.**
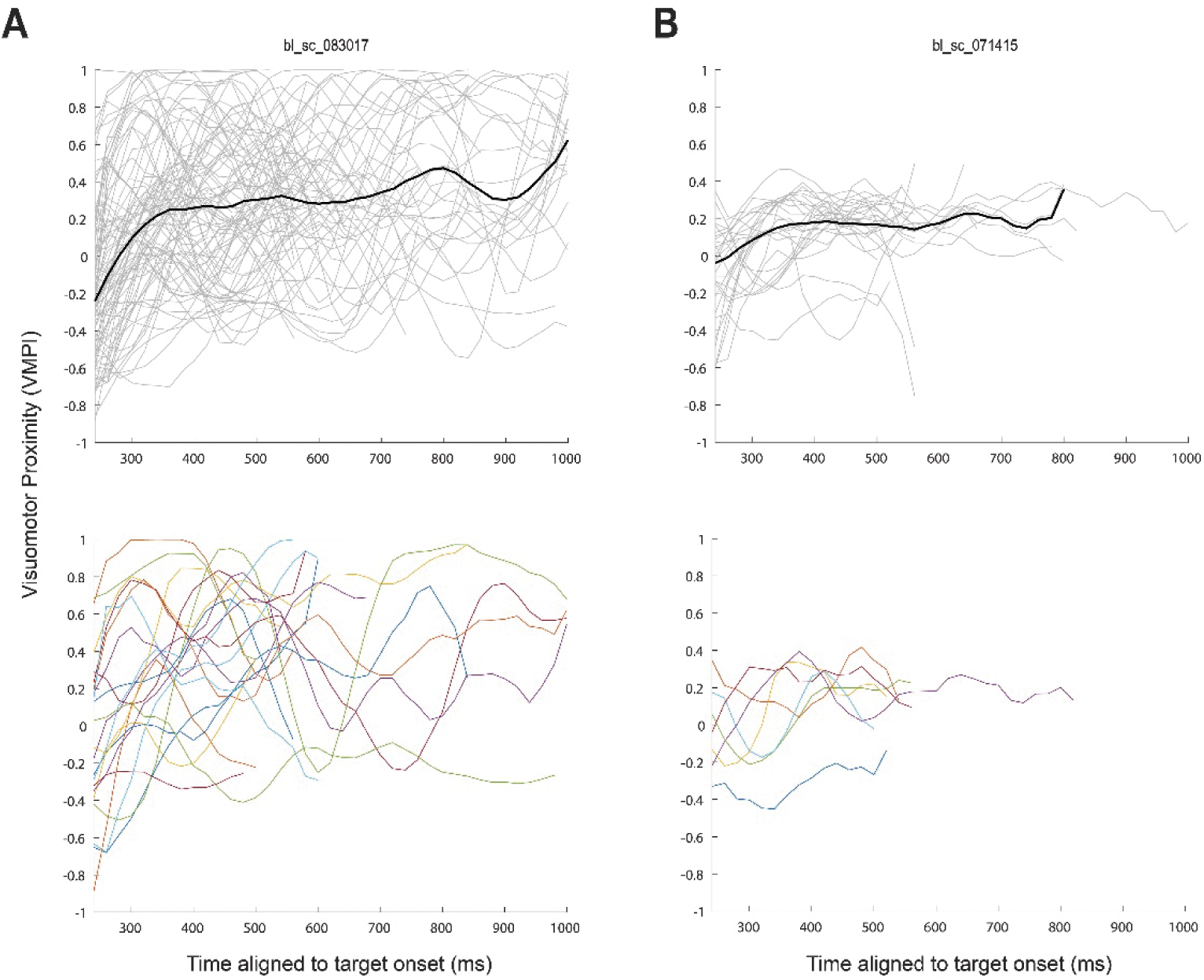
Single-trial delay period VMPI dynamics are highly variable. **A.** Top – VMPI values on individual trials (gray) of an example session (same as Figures 3, 5, and 7) after removing trials in which a microsaccade was made during the delay period. The across-trial median trace is shown in black for all times with at least ten data points (i.e., trials). All traces have been smoothed with a 5-point moving average filter. Bottom – Individual trials from the same example session, subsampled from the entire pool of no-microsaccade trials and individually colored to highlight the across-trial variability in the VMPI trace. **B.** Same as (A) but for a second example session. Here, the range of VMPI values around the across-trial median is much smaller, yet a similarly broad range of dynamics is observable in single trials.

**Supplementary Figure 3.**
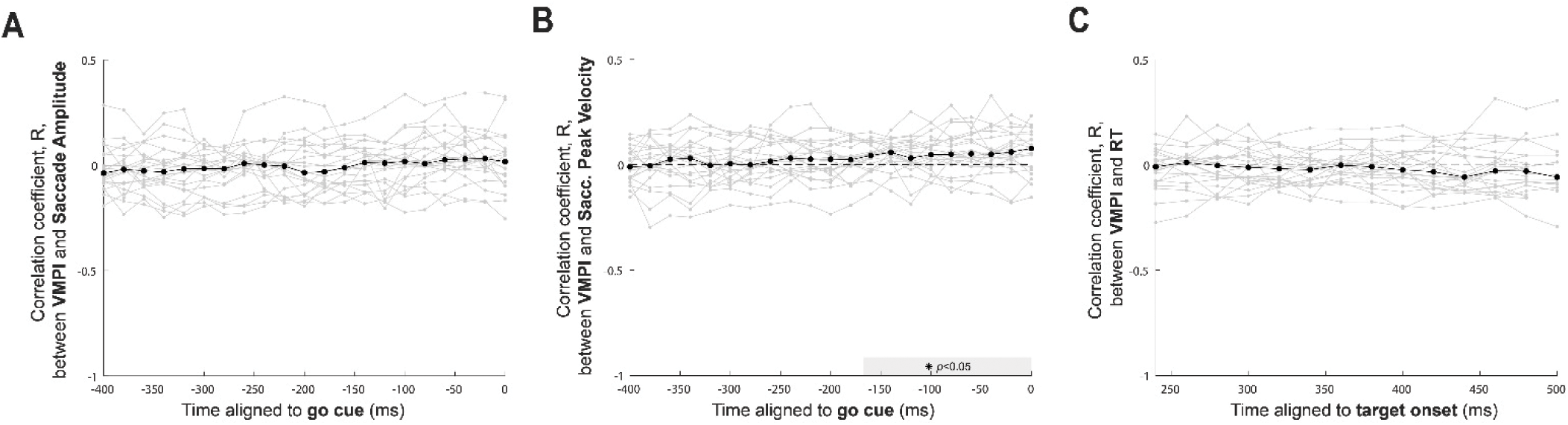
The relationship between VMPI and saccade metrics is variable. **A.** Across-session median correlation coefficient, R, between VMPI and saccade amplitude, for times leading up to the go cue. Same conventions as in Figure 8. There is never a significant correlation between the two variables (Wilcoxon signed rank test). **B.** Same as in (A) but for correlations between VMPI and single-trial peak velocity of the saccade. Time bins in which the median correlation coefficients were significantly different from zero (p<0.05, Wilcoxon signed rank test) are shaded along the x axis in gray. A positive relationship between the two variables emerges approximately 160 ms before the go cue. **C.** Same as in Figure 8A but for correlations between the VMPI value aligned to the beginning of the delay period and single-trial reaction time (RT). No significant correlations are observed (one-tailed Wilcoxon signed rank test).

**Supplementary Figure 4.**
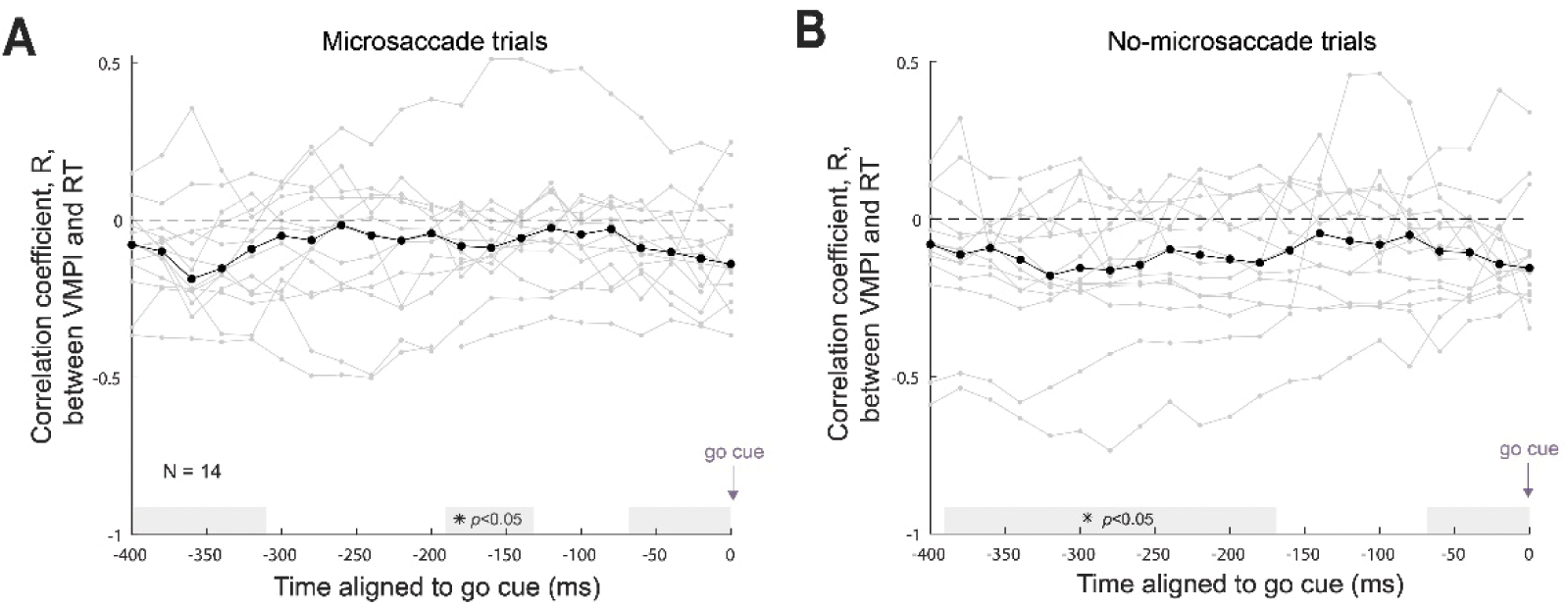
VMPI is correlated with RT for subsets of trials with and without microsaccades. The VMPI value is significantly correlated with saccadic reaction time (RT) at many time points leading up to each trial’s go cue time (p<0.05, Wilcoxon signed rank test) for the subsets of trials in which either a microsaccade (**A**) or no microsaccade (**B**) was made. Same conventions as in Figure 8, with the across-session median correlation coefficients shown in black and individual sessions’ correlation values shown in gray. The relationship between VMPI and RT qualitatively persists in both subsets of trials despite the underpowered analysis due to fewer trials and fewer datasets.

**Supplementary Figure 5.**
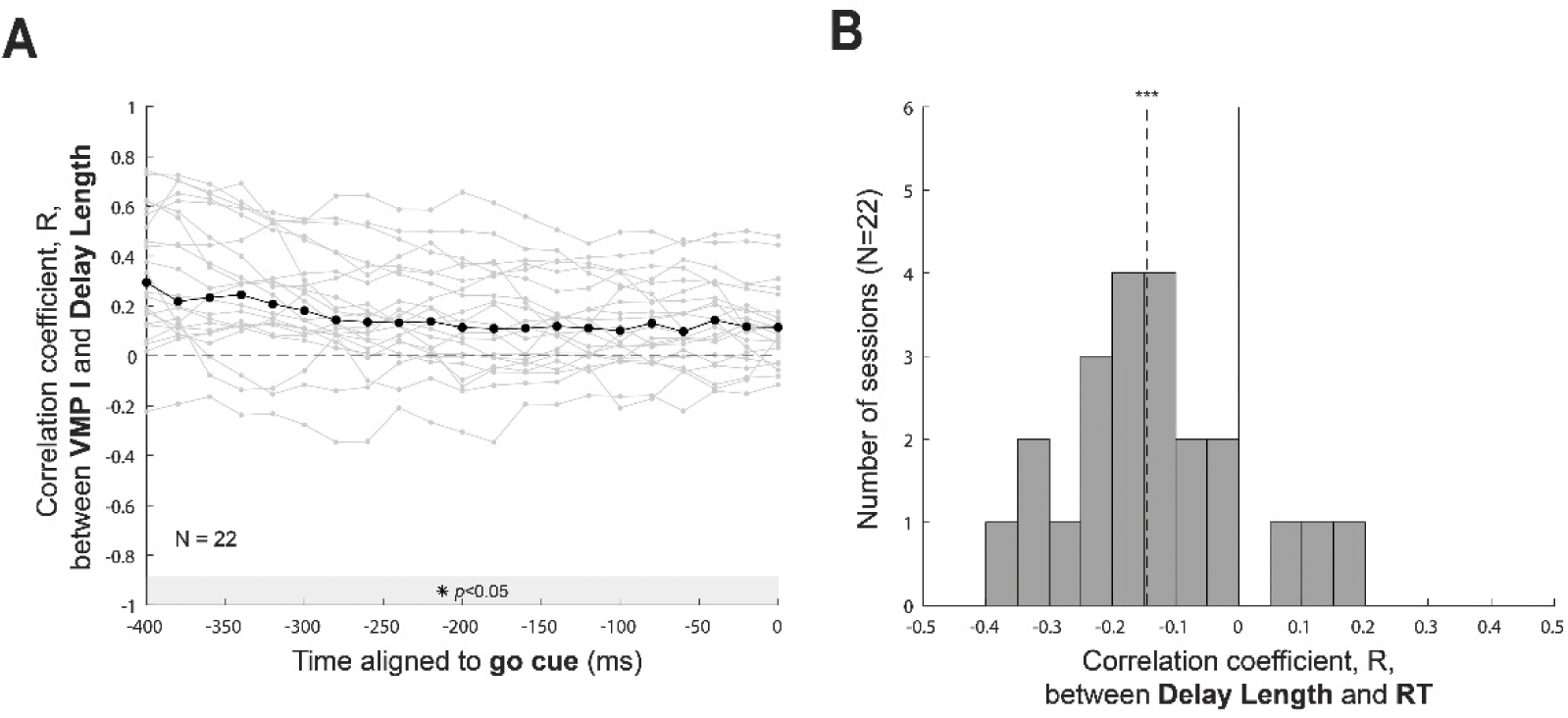
Delay lengths are correlated with both VMPI and RT. **A.** The VMPI value is significantly correlated with the delay period length even 400ms before each trial’s go cue time (p<0.05, Wilcoxon signed rank test). Same conventions as in Figure 8 and Supplementary Figure 4, with the across-session median correlation coefficients shown in black and individual sessions’ correlation values shown in gray. **B.** Histogram of correlation values between each trial’s delay period length and saccadic reaction time (RT) for the 22 sessions included in analysis of sensorimotor transformation. The across-session median correlation coefficient (−0.145) was significantly less than zero (p<0.001, one-tailed Wilcoxon signed rank test), indicating an inverse relationship between delay period length and reaction time.

**Supplementary Figure 6.**
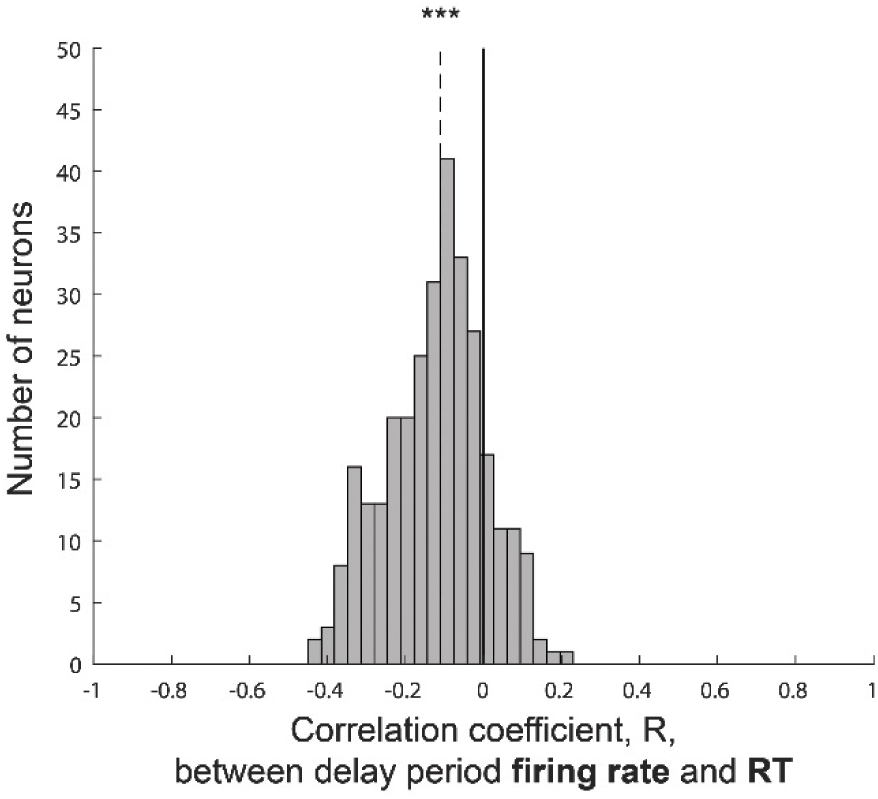
Single-neuron firing rates are also correlated with RT. Histogram of correlation values between single-neuron average firing rate across the full delay period of each trial and saccadic reaction time on that trial for all sorted neurons across all datasets. The across-session median correlation coefficient (−0.110) was significantly less than zero (p<0.001, one-tailed Wilcoxon signed rank test), indicating an inverse relationship between average firing rate across each trial’s delay period and reaction time.

